# Preneoplastic lesions fimbria pan-proteomic studies establish the fimbriectomy benefit for BRCA1/2 patients and identify early diagnosis markers of HGSC

**DOI:** 10.1101/2020.10.04.325100

**Authors:** Maxence Wisztorski, Philippe Saudemont, Soulaimane Aboulouard, Tristan Cardon, Fabrice Narducci, Yves-Marie Robin, Anne-Sophie Lemaire, Delphine Bertin, Firas Kobeissy, Eric Leblanc, Isabelle Fournier, Michel Salzet

## Abstract

Ovarian cancer is the leading cause of death from gynecologic cancer worldwide; however, the origin of ovarian tumors, particularly for high-grade serous carcinoma (HGSC), is still debated. Accumulated evidence converges towards the involvement of the preneoplastic lesions observed in the fimbriated end of the fallopian tubes. In this study, we propose to carry out an in-depth proteomics analysis of these epithelial lesions (p53 signature, serous tubal intraepithelial carcinoma-STIC and serous tubal intraepithelial lesions-STIL) based on spatially resolved proteomic guided by IHC technique. We identified specific clusters related to each preneoplastic lesions, specific protein mutations based on Cosmic database and a Ghost proteome translated from non-coding RNAs and alternative ORFs, using the OpenProt database. Protein networks have been constructed from each cluster utilizing systems biology platform. Generated data were used to confirm the potentially dormant character of the STIL lesion and the more aggressive profile of the STIC which appears closer to HGSC than other lesions. In summary, our results established the chronological mechanisms and genesis of different ovarian cancer phenotypes but also identified the early diagnostic markers of HCSC guiding an adapted therapy and a better patient care.

## INTRODUCTION

Whereas the ovarian cancer (OC) is only at the seventh position in women’s cancer, it is by far the most lethal (only 30-40% survival rate is shown within five years). Amongst the different classes, HGSC is the most represented (>75%) and within this class, 10% of the patients are mutated for *BRCA1/2* or present with Lynch syndrome [1–3]. These patients show a 20-40 percent increased risk of developing HGSC between 30 and 35 years old (versus 1.4% in the global population). During the last 10 years, special attention has been geared towards pathological examination of operative specimens of early OC and prophylactic bilateral adnexectomy performed in women at risk of OC due to an inherited mutation in *BRCA* genes. Most of ovarian carcinomas related to BRCA 1 or 2 mutations are of fallopian tube origin and especially from its distant part called the fimbria. These tubal, ovarian or primary peritoneal carcinomas are quite always of high-grade serous type. They cannot be effectively screened due to the quickness of their evolution. In this context, a laparoscopic bilateral salpingo-oophorectomy (BSO) is the recommended prophylactic procedure. Some BRCA-mutated women (or at-risk of ovarian cancer but without identified mutation) are reluctant to undergo this procedure considering the numerous adverse effects on body and quality of life, especially when hormonal replacement is forbidden. This refusal makes them at risk of developing a serous pelvic carcinoma. The goal of the bilateral laparoscopic radical fimbriectomy we suggest, is to suppress the tubal source of possible dysplastic cells from which can stem this high-grade tumor, while preserving a natural ovarian hormonal secretion. This procedure offered only in case of rejection of BSO intends to salvage these high-risk women, and keep them under an adapted surveillance of their gonads. The reduction in benefits of castration over breast cancer risk, should have been addressed and accepted by women with this specific history. Complementary castration is recommended at 50 years old This new prophylactic operation, based on recent information on ovarian carcinogenesis, preserves femininity and possibly fertility as well. It seems an interesting option for selected women who are reluctant to undergo prophylactic BSO. Its efficacy needs to be assessed. Our work will evaluate Müllerian cells in the fimbria end of the fallopian tube to find a relationship between the pre-neoplastic lesions and high-grade ovarian serous carcinomas (HGSC), a type II ovarian carcinoma. Based on Sectioning and Extensively Examining the Fimbriated End Protocol (SEE-FIM) [4], a systematic serial examination of the fallopian tubes is performed, coupled with IHC evaluation of p53 and Ki-67 expression. In this group of patients, an unusual rate of certain cancer or at least cellular abnormalities were observed in the fallopian tube epithelium (FTE), especially at its terminal end, the fimbria [5, 6]. Three main lesions are defined according to the IHC pattern for the two markers and the architectural alterations of the cells. The p53 signature is identified by a small number of epithelial cells (10-20) with a proliferation activity like the adjacent normal epithelium, but with a p53 staining pattern corresponding to a missense *TP53* mutation. STIC (serous tubal intraepithelial carcinoma) involves many cells with architectural and nuclear alterations, *TP53* mutations and a high proliferative activity. STILs (serous tubal intraepithelial lesions) is characterized by a lower level of abnormalities compared to the STIC along with a normal proliferative activity [7, 8]. Series of morphological changes concomitant with multi-step accumulation of molecular and genetic alterations on pre-neoplastic lesions of FTE hypothesized that HGSCs derive their origin, not in the ovary, but in the fimbria part of the fallopian tube [9–11]. In fact PAX8, a Mullerian marker has been found to be expressed in most HGSC but not calretinin (mesothelial marker) [4, 12, 13]. This view has not been universally accepted, primarily as it conflicts with traditional theories on the origins of OC, and secondarily owing to variation in the detection of tubal lesions in association with HGSC, which may in turn be due to differences in sampling or difficulties in diagnostic interpretation [14, 15]. Recent studies have provided a stepwise progression of FTE to precursor lesions to carcinoma, with the aid of L p53 signature L STIL - STIC - HGSC sequence’s model [11, 16, 17] .

In this context, we propose to use the state-of-the-art of the spatially resolved proteomics as we previously applied on other cancers [18–22]. However, this work has a novel component by using a pathologist guided IHC slide of formalin fixed paraffin embedded (FFPE) tissue section of patients presented p53 signature lesion, STIL lesions and STIC lesions which will be compared to non-pathological fallopian tissues and HGSC. Here, we investigate the different pre-neoplastic lesions, their proteome, the mutations detected in proteins, the proteome translated from alternative ORF, so called the Ghost proteome and finally construct proteome pathways indicative of underlying mechanisms [23–26].

We sought to exploit the benefits of this innovative strategy to improve our understanding of the different molecular events across the pre-neoplastic lesions, their progression, and their possible link with ovarian HGSC.

## EXPERIMENTAL PROCEDURES

### Experimental Design and Statistical Rationale

Shotgun proteomics experiments formalin fixed paraffin embedded (FFPE) tissue section of patients presented p53 signature lesion, STIL lesions and STIC lesions which will be compared to non-pathological fallopian tissues and HGSC were conducted in biological triplicate.

Statistical analysis: For the proteomics statistical analysis of extracted proteins or secreted media, only proteins presenting as significant by the ANOVA test were used with FDR 5%. Normalization was achieved using a Z-score with a matrix access by rows. Obtained data from Western Blot were reported as mean ± SEM. Mean values among different experimental groups were statistically compared by one-way ANOVA tests using Graph pad PRISM software or by student t-test.

### Reagents and Chemicals

For the different experiments, we used high purity chemicals from various suppliers:

HPLC grade methanol (MeOH), ethanol (EtOH), Chloroform (CHCl3), acetonitrile (ACN), water and AR grade trifluoroacetic acid (TFA) were obtained from Thermo Fisher Scientific (Courtabœuf, France). Trifluoracetic acid (TFA, 99%), and ammonium bicarbonate (NH4HCO3) were purchased Sigma-Aldrich (Saint-Quentin Fallavier, France). Xylene and formic acid (FA, ≥96%) are from Biosolve (Dieuze, France). Sequencing grade modified porcine trypsin was obtained from Promega (Charbonnieres, France).

### Case selection and sample processing

Tissues were obtained from patients of the Centre Oscar Lambret (Lille, France. All experiments were approved by the local Ethics Committee (CPP Nord Ouest IV 12/10) in accordance with the French and European legislation. Prior to the experiments, patients signed an informed consent and authorization form describing the experimental protocol. No personal information was used in these experiments, and a random number was assigned to each sample. In our study, all samples were FFPE tissue from prophylactic adnexectomies of women with BRCA mutations. Our study was performed on patients presenting pre-neoplastic lesions without any concomitant ovarian high-grade serous carcinoma. All samples are examined following SEE-FIM (Sectioning and Extensively Examining the FIMbria) protocol [27]. On these samples, tissues sections of 7 µm thickness are done using a microtome and were deposited on glass slide, HPS (Haematoxylin Phloxine Saffron) and immunostaining against P53 and Ki-67 (Dako, Japan) are realized and examined by a pathologist to find preneoplastic lesions (p53/STIL/STIC/HGSC) [28]. A cohort of eight patients were selected. Four patients presented p53 signature lesion, 2 patients with STIL lesions, 2 patients with STIC lesions, 1 with HGSC (**Table 1**). The normal tissue was analyzed from normal part (presenting no abnormalities at IHC) of the tissue section from patients with p53 signature lesion.

**Table 1:**
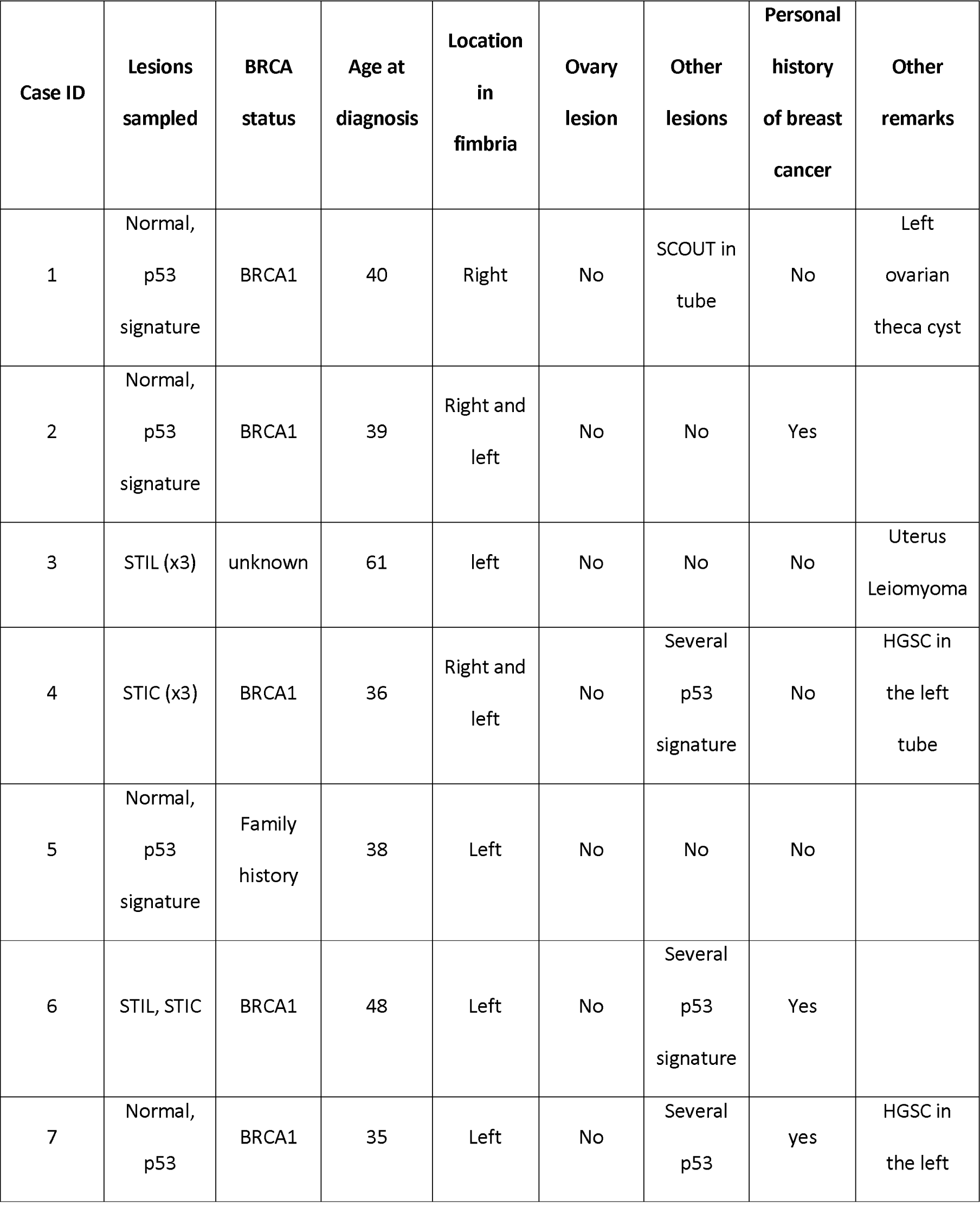
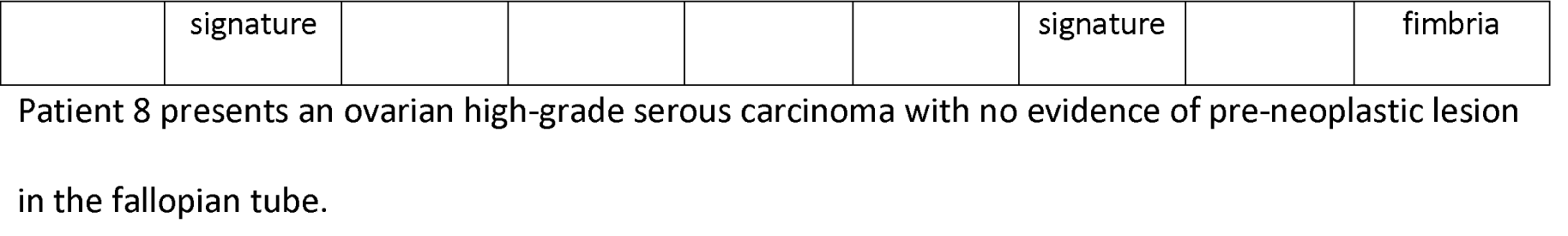
Description of clinical characteristics of patients with preneoplastic lesion in the fallopian tube fimbria.

| Case ID | Lesions sampled | BRCA status | Age at diagnosis | Location in fimbria | Ovary lesion | Other lesions | Personal history of breast cancer | Other remarks |
| --- | --- | --- | --- | --- | --- | --- | --- | --- |
| 1 | Normal, p53 signature | BRCA1 | 40 | Right | No | SCOUT in tube | No | Left ovarian theca cyst |
| 2 | Normal, p53 signature | BRCA1 | 39 | Right and left | No | No | Yes |  |
| 3 | STIL (x3) | unknown | 61 | left | No | No | No | Uterus Leiomyoma |
| 4 | STIC (x3) | BRCA1 | 36 | Right and left | No | Several p53 signature | No | HGSC in the left tube |
| 5 | Normal, p53 signature | Family history | 38 | Left | No | No | No |  |
| 6 | STIL, STIC | BRCA1 | 48 | Left | No | Several p53 signature | Yes |  |
| 7 | Normal, p53 | BRCA1 | 35 | Left | No | Several p53 | yes | HGSC in the left |
|  | signature |  |  |  |  | signature |  | fimbria |

Patient 8 presents an ovarian high-grade serous carcinoma with no evidence of pre-neoplastic lesion in the fallopian tube.

### On-tissue spatially resolved proteomics

Complete workflow for sample preparation is illustrated in Figure 1A

**Figure 1:**
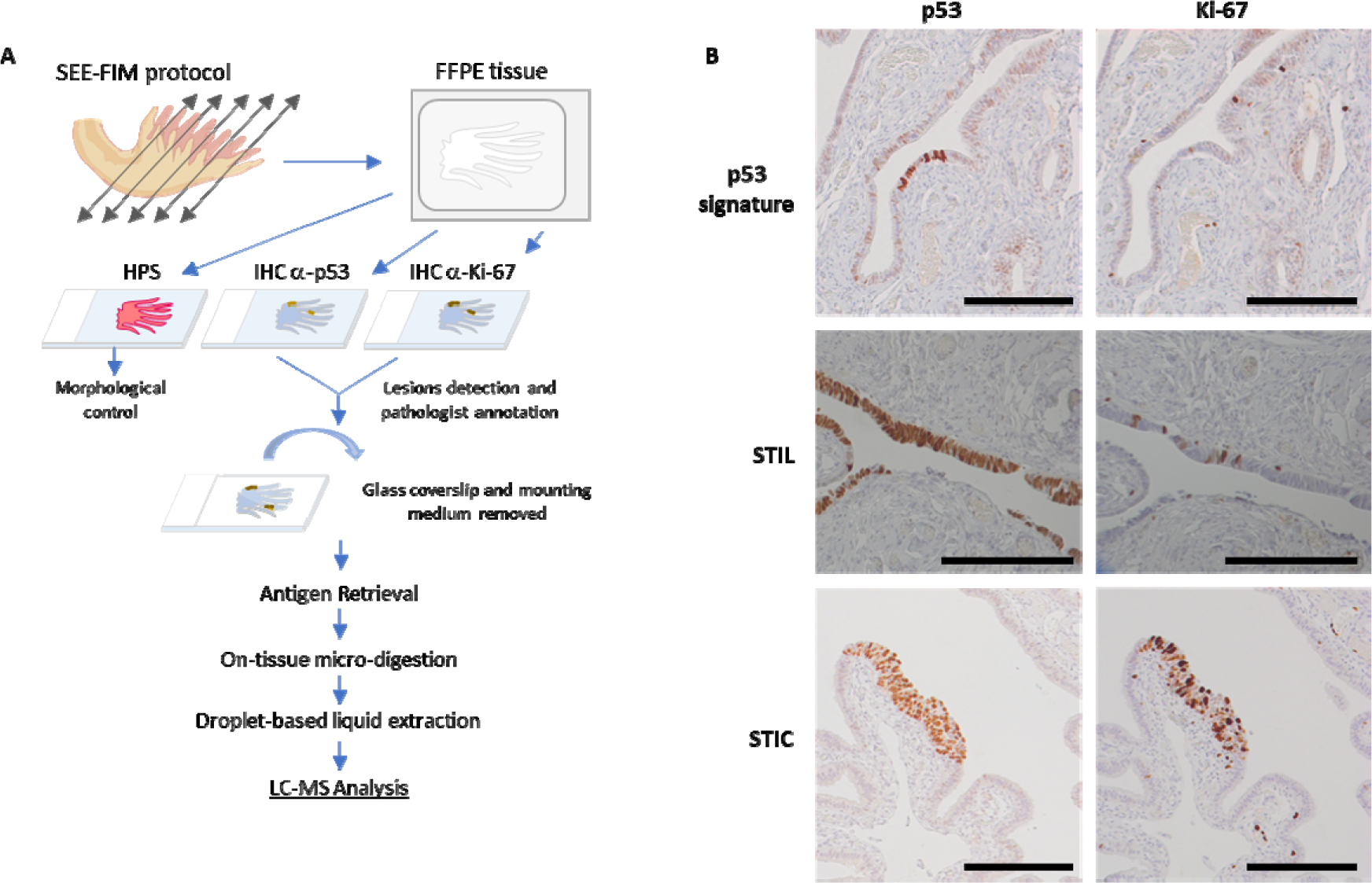
Workflow for spatially resolved proteomic using IHC tissue section. **A)** Protocol based on tissue from SEE-FIM protocol. After IHC against p53 and Ki-67, the coverslip glass and the mounting medium are removed to access the tissue section. A digestion of the lesion is performed directly on the tissue section and a droplet-based liquid extraction is performed to recover the peptides prior to MS-based proteomics analysis. **B)** Pre-neoplastic lesions found in the fallopian tube are defined by p53 positivity and Ki-67 index. For p53 signature: accumulation of p53 in at least 12 cells without morphological abnormalities and low Ki-67 index; STIL: same accumulation of p53 in more than 20 cells with some morphological abnormalities and an higher Ki-67 proliferating index (10-40%); STIC: high p53 and Ki-67 index and cells atypical morphology (carcinoma-like).

### Sample preparation

Same slides as the one used and annotated by the pathologist were unmounted, and the resin removed by soaking them overnight in xylene and rinsing them with xylene and ethanol baths. The tissues are rehydrated using 5’ each successive bath of decreasing ethanol degree (2x95°, 1x30°) and two baths of 10mM NH4HCO3 buffer. Then, an antigen retrieval step was performed to increase the trypsin access to biomolecules. For this, the slides are dipped in 90°C pH9 20mM Tris for 30 minutes, rinsed twice with NH4HCO3 and dried under vacuum at room temperature.

### *In situ* trypsin digestion

On localized pre-neoplastic lesions thanks to the immunostaining, tryptic digestion was performed using a Chemical Inkjet Printer (CHIP-1000, Shimadzu, Kyoto, Japan). The trypsin solution (40µg/mL, 50mM NH4HCO3 buffer) was deposited on a region defined to 1mm² during 2h. During this time, the trypsin was changed every half-hour. With 350 cycles and 450 pL per spot, a total of 6.3µg of trypsin was deposited. To stop digestion, 0.1% TFA was spotted during 25 cycles.

### Liquid Extraction

After microdigestion, the spot content was gathered by liquid microjunction using the TriVersa Nanomate (Advion Biosciences Inc., Ithaca, NY, USA) using Liquid Extraction and Surface Analysis (LESA) settings. With 3 different solvents mixture composed of 0.1% TFA, ACN/0.1% TFA (8:2, v/v), and MeOH/0.1% TFA (7:3, v/v). A complete LESA sequence run 2 cycles for each mixture composed of an aspiration (2µL), a mixing onto the tissue, and a dispensing into low-binding tubes. For each interesting spot, 2 sequences are pooled into the same vial.

### MS-based proteomic NanoLC-ESI-MS^2^

After the liquid extraction, samples were freeze-dried in a SpeedVac concentrator (SPD131DPA, Thermo Scientific, Waltham, Massachusetts, USA), reconstituted with 10µL 0.1% TFA and subjected to solid phase extraction to remove salts and concentrate the peptides. This was done using a C-18 Ziptip (Millipore, Saint-Quentin-en-Yvelines, France), eluted by an ACN/0.1% TFA (8:2, v/v) and then the samples were dried for storage. Before analysis, samples were suspended in 20µL ACN/0.1% FA (2:98, v/v), deposited in vials and 10µLs were injected for analysis. The separation prior to MS used online reversed-phase chromatography realized with a Proxeon Easy-nLC-1000 system (Thermo Scientific) equipped with an Acclaim PepMap trap column (75 μm ID x 2 cm, Thermo Scientific) and C18 packed tip Acclaim PepMap RSLC column (75 μm ID x 50 cm, Thermo Scientific). Peptides were separated using an increasing amount of acetonitrile (5%-40% over 140 minutes) and a flow rate of 300 nL/min. The LC eluent was electrosprayed directly from the analytical column and a voltage of 2 kV was applied via the liquid junction of the nanospray source. The chromatography system was coupled to a Thermo Scientific Q-Exactive mass spectrometer. The mass spectrometer was programmed to acquire in a data-dependent mode for the 10 most intense peaks. The survey scans were acquired in the Orbitrap mass analyzer operated at 70,000 (FWHM) resolving power. A mass range of 200 to 2000 m/z and a target of 3E6 ions were used for the survey scans. The MSMS analysis was performed using HCD with a normalized collision energy of 30 eV, a mass range between 200 to 2000, an AGC of 5e4 ions, a maximum injection time of 60ms and resolution set at 17,500 FWHM. The method was set to analyze the top 10 most intense ions from the survey scan and a dynamic exclusion was enabled for 20 s.

### Data Analyses

MS/MS spectra were processed using MaxQuant (version 1.5.6.5, [29, 30]). Peptides identification were obtained according to a target decoy searches against Homo sapiens Uniprot database (version of 2017_02, 20,172 sequences) and database containing 262 commonly detected contaminants. The human uniport database was used as forward database and a reverse one for the decoy search was automatically generated in MaxQuant Mass tolerances were set to 10 ppm and 20 ppm respectively for parent and fragments measurement. Enzyme specificity was set to “trypsin” with a maximum of 2 missed cleavages allowed. Methionine oxidation and acetylation of protein N-terminal were set as variable modifications. FDR < 1% was set for peptides and proteins identification. Two peptides with one unique were necessary to assess protein identification. For label-free quantification, the MaxLFQ algorithm was used [31]. The option “Match Between Runs” was enabled to maximize the number of quantification events across samples. This option allowed the quantification of high-resolution MS1 features not identified in each single measurement. Data generated by MaxQuant were analyzed using Perseus (version 1.6.2.3, [32]. LFQ values were used and proteins were removed if found in the category only identified by site modifications, in the decoy reverse database or identify in the contaminant database. We removed proteins that were not presented in at least three of four replicates. We took the average expression per group to perform a comparison and visualized using a Venn diagram. Individual LFQ values were used to perform a multi scatter plot and calculate a Pearson Correlation between samples. Missing values were imputed based on normal distribution (width=0.3, down-shift= 1.8). Principal component analysis (PCA) was done to compare the protein content of each sample. An ANOVA Multi sample test was done and consolidated by a Permutation-based FDR (FDR< 0.05, 250 randomizations). A specific comparison between normal and p53 signature samples was performed using a student’s T-test.

Proteins with significant differences were filtered out and values were z-scored. The samples were then clustered according to a Euclidean average as a distance measure for column and row clustering. The MS data sets and Perseus result files used for analysis were deposited at the ProteomeXchange Consortium [33] (http://proteomecentral.proteomexchange.org) via the PRIDE partner repository [33] with the data set identifier PXD020024 (for reviewer access only, Username; Password: x88jqaDV).

Up-regulated and down-regulated proteins in the different groups were used to perform an annotation analysis of gene ontology terms by using Funrich (v3.1.3) [34]. A hypergeometric test was performed against all annotated gene/protein list by comparing the multiple datasets. Enrichment for biological process, transcription factor and cellular component were present as a bar chart. PANTHER Classification System (v14.1) was also used. PANTHER Overrepresentation test (Released 20190701) was performed using each list of up- or down-regulated proteins as “analyzed list” and Homo sapiens as “reference list”. Fisher’s Exact test with false discovery rate correction was used.

Subnetwork Enrichment Analysis (SNEA) from Elsevier’s Pathway Studio version 10.0 (Elsevier) [35, 36] was used to extract statistically significant altered biological and functional pathways in the different clusters of proteins.

The immunohistochemical (IHC) data of different proteins of interests were investigated from the Human Protein Atlas (HPA) database (http://www.proteinatlas.org, [37]). Evaluation of the prognostic effects of these proteins on the overall survival (OS) were also extracted from HPA. Comparisons are performed with the 20 genes of highest significance associated with unfavorable prognosis for ovarian cancer, cervix cancer, endometrial cancer, and breast cancer.

### Mutation identification

The MS data were also processed to search for potential mutated peptides using the XMAn database [38]. This database contains information on mutations observed in cancers and diseases. Proteome Discoverer 2.1 was used to query the data using MS Amanda as a search node against the XMAn database, the human Uniprot database, and a database containing potential contaminants. Peptides identified only in the XMAn database at a high level of confidence were selected, and the MSMS spectra were manually inspected to confirm the presence of the mutation.

### Alternative Proteins identification

RAW data obtained by nanoLC-MS/MS analysis were analyzed with Proteome Discoverer V2.3 (Thermo Scientific) using the Label Free Quantification node, the protein database is downloaded from Openprot (https://openprot.org/)[39]. This database included RefProt, novel isoforms and AltProts predicted from both Ensembl and RefSeq annotations (Ensembl: GRCh38.83, RefSeq: GRCh38p7) for a total of 658,263 entries. The following processing and consensus parameters are used with: Trypsin as an enzyme, 2 missed cleavages, methionine oxidation as variable modification, precursor mass tolerance: 10 ppm and fragment mass tolerance: 0.1 Da. The validation was performed using Percolator with a protein strict FDR set to 0.001. A consensus workflow was then applied for the statistical arrangement, using the high confidence protein identification. Results are filtered to keep master protein and high confidence protein FDR. Then results are extract in table, in order to use the LFQ values in PERSEUS, where an ANOVA statistical test is performed and the results represented by a diagram, like heatmap has described above.

## RESULTS

### Compatibility between IHC and droplet-based liquid extraction for spatially resolved proteomics

A cohort of 8 patients presenting *BRCA* mutations or hereditary susceptibility, were selected for the experiments (**Table 1**). In this study, we used immunostained tissue section coming from a SEE-FIM protocol (**Figure 1A**). This protocol of systematic examination of the totality of the fimbria is difficult to perform and will not allow observation of a lot of pre-cancerous lesions. Indeed, the number of samples is then critically limited allowing minimal slide examination. Moreover, the pre-cancerous lesions present an extremely limited number of cells on the epithelial layer with no specific morphological features (**Figure 1B**). The presence of p53 signature, STIL and STIC lesions were thus, confirmed-thanks to a double IHC against p53 and Ki-67 markers (**Figure 1B**). In order to process these samples for spatially resolved proteomics, we developed a new protocol directly using the same IHC slide used by the pathologist which has served to the detection of the lesions. The cover slide and the mounting medium are removed by extensive bathing of xylene and ethanol. A visual inspection is carried out to check whether additional washing is required to remove all the mounting medium from the tissue section. One critical point was the antigen retrieval procedure that needs to be performed after tissue rehydration. Next, the region of interest corresponding to a lesion is covered with a droplet of trypsin solution. The resulting tryptic peptides are extracted from the tissue by a droplet-based liquid microextraction derived from the LESA method [19-21, 40-42] and analyzed by MS-based proteomics. Considering all the pre-neoplastic lesions and non-pathological tissues, 10375 unique peptide sequences corresponding to 1617 distinct protein groups have been identified using our workflow. MaxLFQ algorithm was used to perform label free quantification of proteins and resulted in a total of 1571 protein quantifications. After filtering one according to a minimum number of values (2/3 of valid values) in at least one group of the four defined groups, 1046 proteins were obtained. This result is like those obtained in our previous experiments on FFPE tissue.

Furthermore, using immunostained tissue after removing the coverslip glass and the mounting medium does not appear to affect not affect the microproteomic analysis.

To confirm this, we performed correlation analysis based on all quantified proteins. Proteomes inside samples from the same group were similar (mean Pearson correlation 0.92) compared with inter-group variation. The most differences were observed between STIC and STIL lesions (mean of 0.84) (**Figure 2A**). This is in accordance with what is generally observed considering the inter-patient variability, and the protocol we used does not appear to alter the proteomic content of the tissue sections. This result allows us to perform a comparative analysis to obtain a specific proteomic signature of the different lesions.

**Figure 2:**
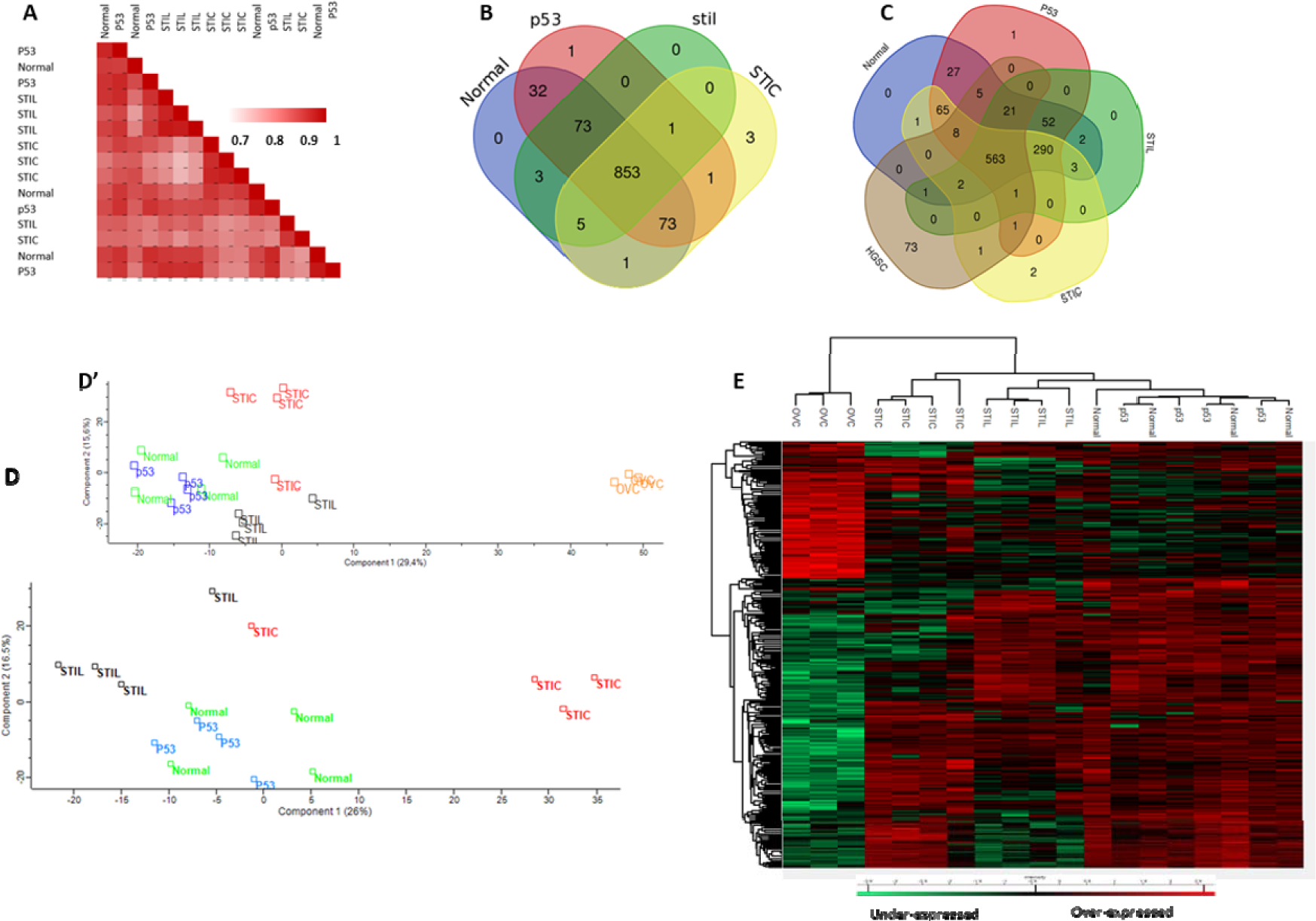
Comparison of the different lesions based on their proteomic signature. **A)** Correlation analysis based on the whole proteome. Ven diagrams showing distribution of identified proteins among **B)** normal tissue and the different lesions and **C)** with addition of HGSC data. Principal component analysis using **D)** normal tissue and the pre-neoplastic lesions and **D’)** with addition to HGSC. E) Hierarchical clustering of the most variable proteins (ANOVA with permutation-based FDR < 0.05) including HGSC data

### Spatially resolved proteomic study of p53 signature, STIL, STIC and normal Fimbria

Comparing the content of each group by averaging the replicates and using a Venn diagram representation, 853 common proteins (81.5% of overlap) between normal tissues and pre-neoplastic lesions were observed (**Figure 2B, Supp. Data 1**). Few unique proteins in each group were observed. One protein is found only in the p53 signature lesion and three exclusives to the STIC lesions. The protein RNA-binding protein 10 (RBM10), found in three of the four p53 signature lesions and not in the normal tissue or other lesions, is an RNA-binding protein located in the nucleus. Human Protein Atlas (HPA) data shows that overexpression of this gene is favorable in terms of prognostic in the case of cervical cancer and endometrial cancer. Three proteins (Glucosamine (N-acetyl)-6-sulfatase (GNS), Upstream binding transcription factor, RNA polymerase I (UBTF) and ATPase family, AAA domain containing 3 (ATAD3A/B)) are exclusives to STIC lesions. Survival data extracted from HPA shows that high expressions of GNS and ATAD3A/B are not favorable for survival in liver cancer. GNS is also poor prognosis for glioma and present the same tendency in ovarian cancer. The expression of UBTF appears to have no impact on survival. Inclusion of HGSC data in the analysis (**Figure 2C, Supp data 2**), revealed that UBTF is present in STIC lesion and in HGSC. Fibrillarin (FBL) is found only in the pre-cancerous lesions and not in the normal tissue. This protein is also observed in HGSC (**Figure 2C**, **Supp. Data 2**). Comparison between the data of the different lesions and the HPA list of the 20 genes of highest significance associated with unfavorable prognosis in ovarian cancer revealed only two proteins (TGFBI and RPL7A), 3 (SUMF2, TGFBI, FASN) with cervix cancer, 1 with endometrial cancer (ASS1) and 4 (PGK1, ATP5F1B, TAGLN2, RAB5C) with breast cancer.

To better understand the modulation registered across the different lesions, principal component analysis (PCA) and hierarchical clustering were performed to visualize the correlation between samples. Principal Component analysis based on whole proteome levels allow visualization of the difference between each group. (**Figure 2D**). Component 1 and 2 account for 42.5% of the total data variation. Samples from normal tissue and p53 signature are closed and could not be easily differentiated. Samples from STIL and STIC lesions differed from normal and p53 signature samples in component 2. STIL lesions are closer to normal/p53 lesions in component 1 than STIC. Adjunction of the HCSG data confirmed the separation of the nature of the proteome between p53 signature, STIL, STIC and HGSC (**Figure 2D’**). Hierarchical clustering showed a significant difference in expression between the 5 groups as shown in the heatmap (**Figure 2E**). Two main branches separate HGSC from the other lesions. The second branch is subdivided again into two subbranches *i.e*., one separating the STIC to p53 signature/Normal/STIL (**Figure 2E**). The second subbranches is then separated between STIL to p53 signature/Normal. The last group (p53/Normal) is difficult to be differentiated. Nevertheless, the difference in cell composition between the ovary and the fallopian tube does not allow a direct conclusion to be drawn on the relationship between HGSC and the pre-neoplastic lesions observed in the fallopian tube.

However, we can observe that the proteomic content of the STIC lesions is closer to HGSC than to the other lesions and that each lesion seems to present a specific proteomic signature.

### Proteomic analyses along the tumoral process

For a deeper analysis of the pre-neoplastic lesions of the FTE, ANOVA was performed without the HGSC data. A total of 197 proteins showed a variation of expression on the four groups (**Supp. Data 3**). A hierarchical cluster based on the expression of these proteins resulted in three groups and two mains clusters (**Figure 3A, Supp. Data 3**). As observed previously in PCA, Normal and p53 signature could not be resolved and formed one group. This group and one composed by the STIL lesions samples clustered together. Proteomics signature of STIC differed from the two previous groups and samples from STIC lesions formed the second cluster suggesting that proteomics profiles differed a lot from the two other groups. The heatmap presents five main clusters of proteins. Proteins that were enriched in STIL and STIC regions were represented in **cluster 1**, those that were more abundant in both normal tissue and p53 signature lesions were represented in **cluster 2**, proteins presented in normal, p53 signature lesion and STIL but not in STIC were represented in **cluster 3**, proteins specifically overexpressed in STIC were represented in **cluster 4** and those specific to normal tissue, p53 signature lesion and STIC were represented in **cluster 5**. **Cluster 1** (STIL/STIC) is composed by only 5 proteins i.e., MARCKS-related protein (MARCKSL1), Myosin-10 (MYH10), Protein SET (SET), Double-stranded RNA-binding staufen homolog-1 (STAU1) and D-3-phosphoglycerate dehydrogenase (PHGDH). The SET protein participates in numerous cellular functions including DNA repair, transcription, cell survival and proliferation. This protein is also involved in many cancer processes such as metastasis, the development of therapeutic drug resistance and plays a key role in tumorigenesis [43]. MARCKSL1 is known to promote the progression of lung adenocarcinoma by regulating Epithelial-mesenchymal transition (EMT) [44]. STAU1 has been previously identified in colorectal cancer [45], whereas PHGDH protein has been identified in ovarian cancer [46, 47]. In **cluster 2** (normal-p53), 15 proteins showed an overexpression for normal and p53 signature lesion. Among these proteins, CD166 antigen (ALCAM/CD166) is shown to be a potential cancer stem cell marker [48]. It has been found as a poor prognosis marker in pancreatic cancer [49] or gastric cancer [50].This protein is known to contribute to local invasion and tumor progression by acting on the detachment of tumor cells [51, 52]. Moreover, TNPO1, ACLY, NME1, FLOT1, KLC4 are proteins already know to be involved in epithelial ovarian cancer [53–55]. Gene Ontology (GO) analyses confirmed that these proteins are involved in cell proliferation, adhesion, and growth (**Figure 3Ba**). **Cluster 3** (normal-p53/STIL) contains 80 proteins. Molecules for the complement and coagulation cascades are present in this cluster (SERPIN A1, SERPIN C1, SERPIN G1, C4a, C9, Vitamin D-binding protein (GC), pro-thrombin (F2), Vitronectin (VTN), kininogen (KNG1)) and 10 members of the collagen family. The PGRMC1 (Membrane-associated progesterone receptor component 1) is known to be involved in ovarian cancer [56] as well as PTGIS (Prostacyclin synthase) [57] and the Protein NDRG1 known to modulate genes involved in ovarian cancer metastasis [58]. GO analyses confirmed that proteins are involved in neoplasia/metastasis, angiogenesis and apoptosis (**Figure 3Bb**). **Cluster 4** (STIC only) contains 36 proteins from which proteins involved in cell migration can be found such like CFL1 (Cofilin-1), PFN1 (profilin-1), PPL (periplakin), FLNB (filamin B), PDLM1(PDZ and LIM domain protein 1), IQGAP1 (Ras GTPase-activating-like protein), MYH14 (Myosin-14) and in metabolism (CS (Citrate synthase), DECR1 (2,4-dienoyl-CoA reductase), GSR (Glutathione reductase), LAP3 (Cytosol aminopeptidase), GLUD1 (Glutamate dehydrogenase 1). Proteins of this cluster are involved in cell differentiation and proliferation and still in neoplasia/metastasis (**Figure 3Bc**). The last cluster (normal-p53/STIC) is composed of 61 proteins. Several proteins involved in exosomes have been identified (Annexins A1, A2, A4, A13, EZR (ezrin), HSPA8 (Heat shock 71kDa protein 1A))., other are involved in stress (PRDX1 ‘Peroxiredoxin-1), HSP8, HSPA1 (Heat shock 70 kDa protein 1), HSPA2 (Heat shock 70 kDa protein 2), ST13 (Hsc70-interacting protein), DNAJB6 (DnaJ homolog subfamily B member 6), VCP (Transitional endoplasmic reticulum ATPase), TUBB4B (Tubulin beta-4B chain)). This cluster shows an overrepresentation of proteins involved in neoplasia and cancer transitions (**Figure 3Bd**). Functional enrichment analyses point out differences between clusters. Different biological processes have been identified *i.e.* immune response, regulation of immune response, cell growth and/or maintenance, metabolism, energy pathways, signal transduction and protein metabolism (**Figure 4A**). Enriched transcription factors (TF) that regulate the overexpressed proteins in the different clusters were also obtained (**Figure 4B**). The cluster containing proteins overexpressed in normal-p53 signature show a high abundance of proteins involved in metabolism and energy pathways as well as an enrichment of SP1, SP4 and TEAD1 transcription factors that targeted the most genes of this cluster. For the cluster normal-p53/STIL, proteins are mostly involved in cell growth and/or maintenance and protein metabolism. Enrichment of the TFs NFIC and EGR1 was observed. Concerning Normal-p53/STIC implicates proteins in cell growth and /or maintenance, Metabolism, energy pathways and protein metabolism for the biological process and KLF7 as TF. For STIL/STIC cluster, enrichment of the TFs ZEB1 and ETS1is observed. STIC cluster presents a strong enrichment of proteins involved in protein metabolism and regulation of immune response and SP1, SP4 and KLF7 as TFs. We also observed a diminution of the proteins involved in immune response from normal-p53 to STIC. Cellular components analyses reflect that all proteins of the 5 clusters are in cytoplasm or exosomes (**Figure 4C**) but with some differences. In fact, in STIL-STIC and STIC we observed a strong enrichment for exosomes whereas p53 signature and STIC involve cytoplasmic proteins. We also observed a high enrichment of cytoskeletal proteins in STIC lesions. Altogether, these analyses reflect some clear transition occurring between Nomal-p53 signature, p53 to STIL and STIL to STIC with a modulation of proteins involved in cell growth and/or maintenance and the different metabolism processes. To confirm these observations, further analyses were carried out, this time grouping together all proteins overexpressed in each lesion, i.e., normal-p53 signature, STIL and STIC (**Figure 4D, E and F and Supp. Data 3**). In STIL, an enrichment of proteins involved in cell growth and/or maintenance (**Figure 4D**), in integrin cell surface interactions (**Figure 4E**), in extracellular matrix (ECM) structural constituent and cell adhesion molecule activity (**Figure 4F**) is observed and the ones in various metabolism and energy pathways (**Figure 4D and E**), catalytic activity (**Figure 4F**) are lower than for the other cellular transition. We also investigated whether the proteins involved in the Warburg effect [59] are activated to compensate for the overall decrease in the classical metabolism processes observed (**Figure S1**). These proteins are not altered overall with a decreasing rather than an increasing trend, which means that no Warburg effect seems to occur. In STIC, proteins involved in metabolism, energy pathways, cell communication, signal transduction (**Figure 4D and E**), calcium ion binding, catalytic and structural molecule activities (**Figure 4F**) are higher than the other transition. In the same time, immune response (**Figure 4D**), integrin interactions (**Figure 4E**), cell adhesion molecule activity and ECM structural constituent (**Figure 4F**) are highly repressed. Statistical overrepresentation test using PANTHER classification system analyses (**Supp. Data 4**) confirmed such transition.

**Figure 3:**
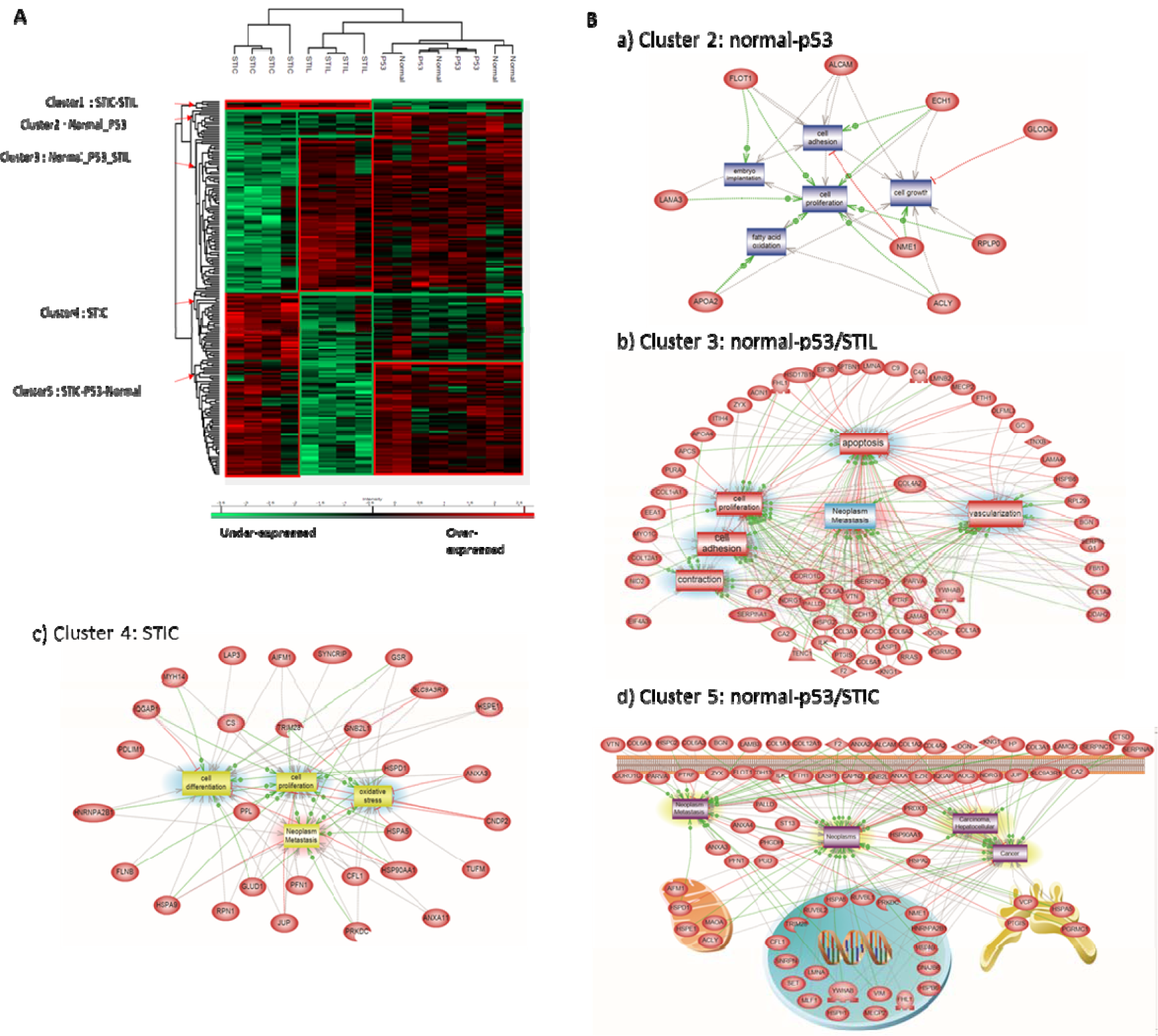
Proteomic analysis of the pre-neoplastic lesions. **A)** Hierarchical clustering of the most variable proteins between normal tissue, p53 signature, STIL and STIC (n=4 for each category, ANOVA with permutation-based FDR < 0.05); **B)** Subnetwork Enrichment Analysis was done to highlight altered biological and functional pathways in the different clusters of proteins.

**Figure 4:**
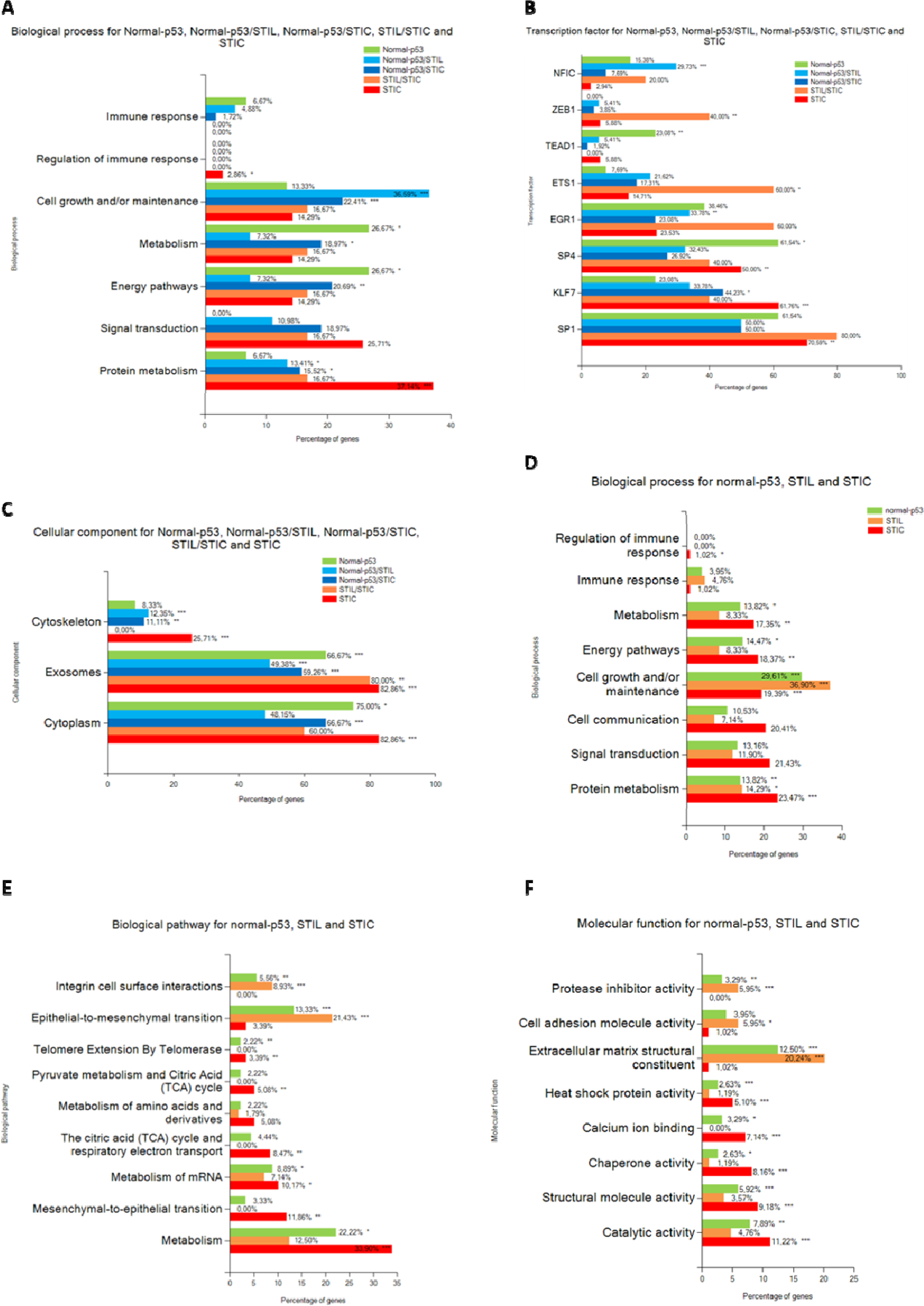
Annotation analysis of gene ontology terms. **A)** Biological process, **B)** Transcription factor and **C)** Cellular component for the clusters of proteins. **D)** Biological process, **E)** Biological pathway and **F)** Molecular function of the proteins overexpressed in each lesion. (hypergeometric test against all annotated gene/protein list of Funrich database, p-value is represented by stars: *** p < 0.001, ** p < 0.01, * p < 0.05 no star for p > 0.05).

Nevertheless, there are still difficulties in distinguishing the differences between p53 signature and normal tissue. Using the Student’s T-test, 34 proteins have been shown to be highly discriminant between Normal and p53 lesions with 26 overexpressed in Normal and 8 in p53 signature (**Figure 5, Supp. Data 5**). **Table 2** presents the immunoreactivity in normal FTE and prognostic effect on the overall survival (OS) in OC for these 8 proteins extracted from the HPA. PTRF, also named CAVIN1, is an unfavorable prognosis marker for ovarian, urothelial, and colorectal cancers, whereas SDPR (known as CAVIN2) is a renal cancer favorable prognosis marker and unfavorable in stomach cancer. The IHC data indicated that these proteins are not detected in normal FTE, show a moderate positivity in OC for CAVIN1 and a strong positivity in a rare case of endometrioid carcinoma of ovary for CAVIN2. SPTAN1, FBLN5 are a favorable prognostic marker in renal cancer but unfavorable in OC (considering a *p* < 0.05). EIF3B, MOB1B and Emilin2 are bad prognosis markers for liver cancer, also for renal cancer for EMILIN-2 and EIF3B and in head neck cancer for EIF3B. Considering the 26 proteins downregulated on p53 signature, different pathways are down represented in these samples, *i.e.,* Neutrophil degranulation, SRP-dependant cotranslational protein targeting to membrane and cadherin binding function.

**Figure 5:**
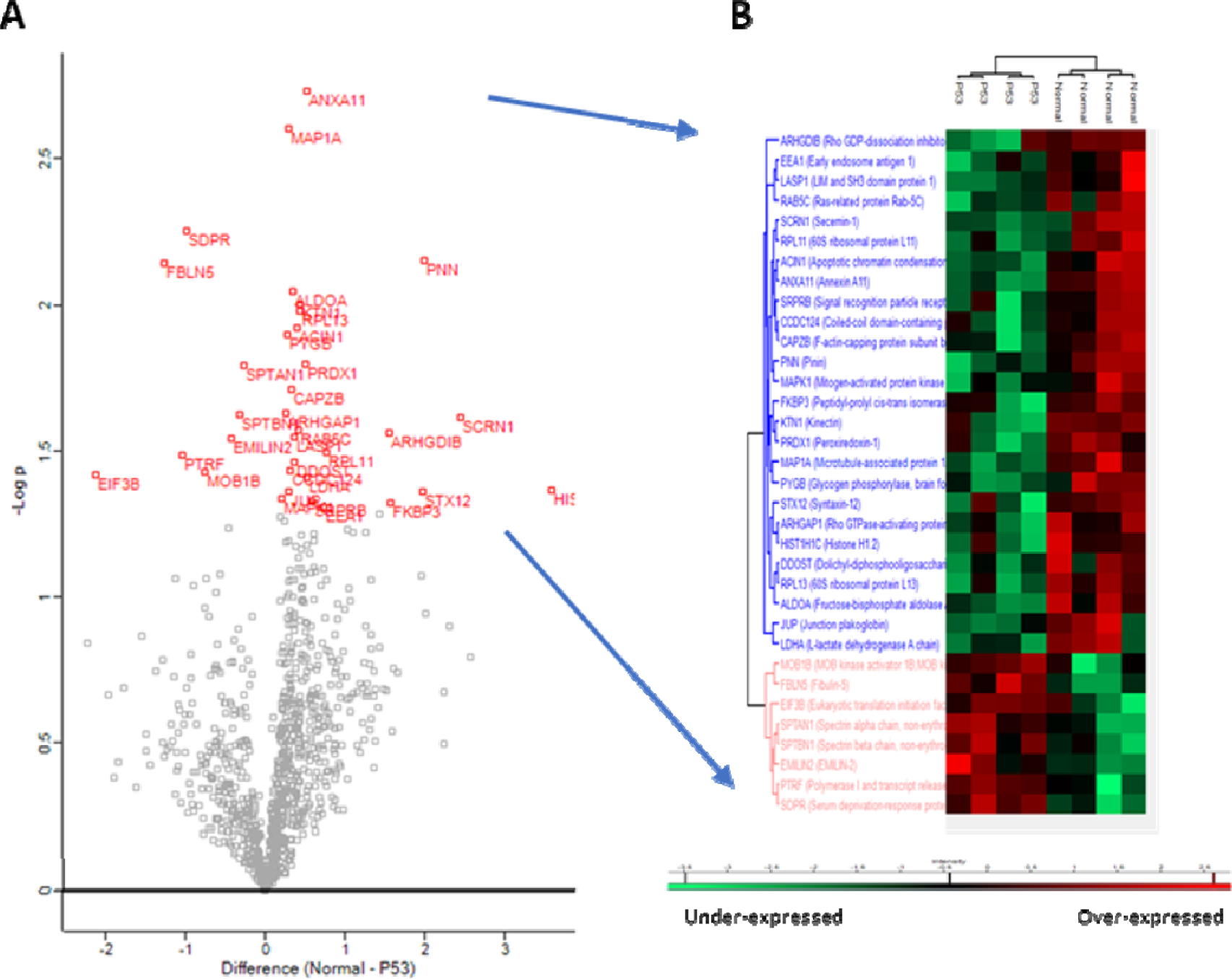
Comparison between p53 signature and normal tissue. A) Visualization of a t-test in form of a volcano plot comparing normal to p53 lesion (proteins with p < 0.05 in red.) B) Hierarchical clustering of the significant variable proteins between normal tissue and p53 signature (t-test with p < 0.05).

**Table 2:** Immunoreactivity and prognostic effect of proteins overexpressed in p53 signature compared to normal tissue. Immunohistochemical staining of proteins in normal FTE tissue and prognostic effect of high expression of the corresponding gene in OVC were extracted from Human protein atlas (data credit: Human Protein Atlas available from v19.proteinatlas.org, p-value is represented by stars *** p < 0.001, ** p < 0.01, * p < 0.05 and no star for p > 0.05)

| Protein name | Immunoreactivity in FTE | Prognostic effect in OVC |
| --- | --- | --- |
| CAVIN1 | Not detected | Unfavorable *** |
| SPTAN1 | Moderate to strong positivity | Unfavorable * |
| FBLN5 | Not detected | Unfavorable * |
| EIF3B | Strong positivity | Not significant |
| MOB1B | Moderate positivity | Not significant |
| EMILIN2 | Moderate positivity | Not significant |
| SPTBN1 | Moderate to strong positivity | Not significant |
| CAVIN2 | Not detected | Not significant |

| **Found in samples** | **Patients\***<br>**Number** | **Protein** | **Mutation** | **Cancers** |
| --- | --- | --- | --- | --- |

### Protein Mutation

Lineage relationship between HGSC and the different lesions are suggested by the presence of same genomics changes and mutations [60]. In order to identify specific modifications that may have occurred during the different potential pre-cancerous lesions, we explored the presence of protein mutations per lesions. For that purpose, we used the human database combined with the XMAn database [61]. This database contains information concerning mutated peptides that could be found in some cancers extracted from the COSMIC database. This database integration resulted in the identification of 83 peptide sequences containing possible mutations (**Supp. Data 6**). For four of them, it was possible to determine the amino acid modification directly in the tandem mass spectra (**Figure 6A**). These peptides are derived from four proteins *i.e.,* the vitamin D binding protein (GC), the polyubiquitin-C (UBC), the Histone H2B (H2B1C) and Histone H3.1 (H31). The mutation found in the protein GC (1296T>G p.D432E), lead the sequence LP**<u>E</u>**ATPTELAK in the protein and. has been found in stomach cancer according to the COSMIC database [62–65]. It was observed in data issued from two patients in a normal tissue and in a p53 signature lesion. Interestingly this protein was found to be significantly under-represented in the STIC lesion (**Supp. Data 3**). Similarly, the mutated peptide of the UBC was found in two normal tissues, two p53 signature lesions and one STIL. The mutationc.368G>C p.G123A, which leads the LIF**<u>A</u>**AKQLEDGR was previously identified in breast cancer The mutated peptide (QVHPDTGIS**<u>T</u>**K) from Histone H2B (mutation c.169T>A p.S57T) is found only in one normal tissue and the protein was not dysregulated (**Supp. Data 3**). This mutation has been identified before in the endometrium. The peptide of the histone H3.1 (VTIMP**<u>R</u>**DIQLAR) was observed in two normal tissue, one p53 signature and one STIL lesion. This has been found to be downregulated in the STIC lesion (**Supp. Data 3**). This mutation (c.368A>G p.K123R) has also been identified in the endometrium.

**Figure 6:**
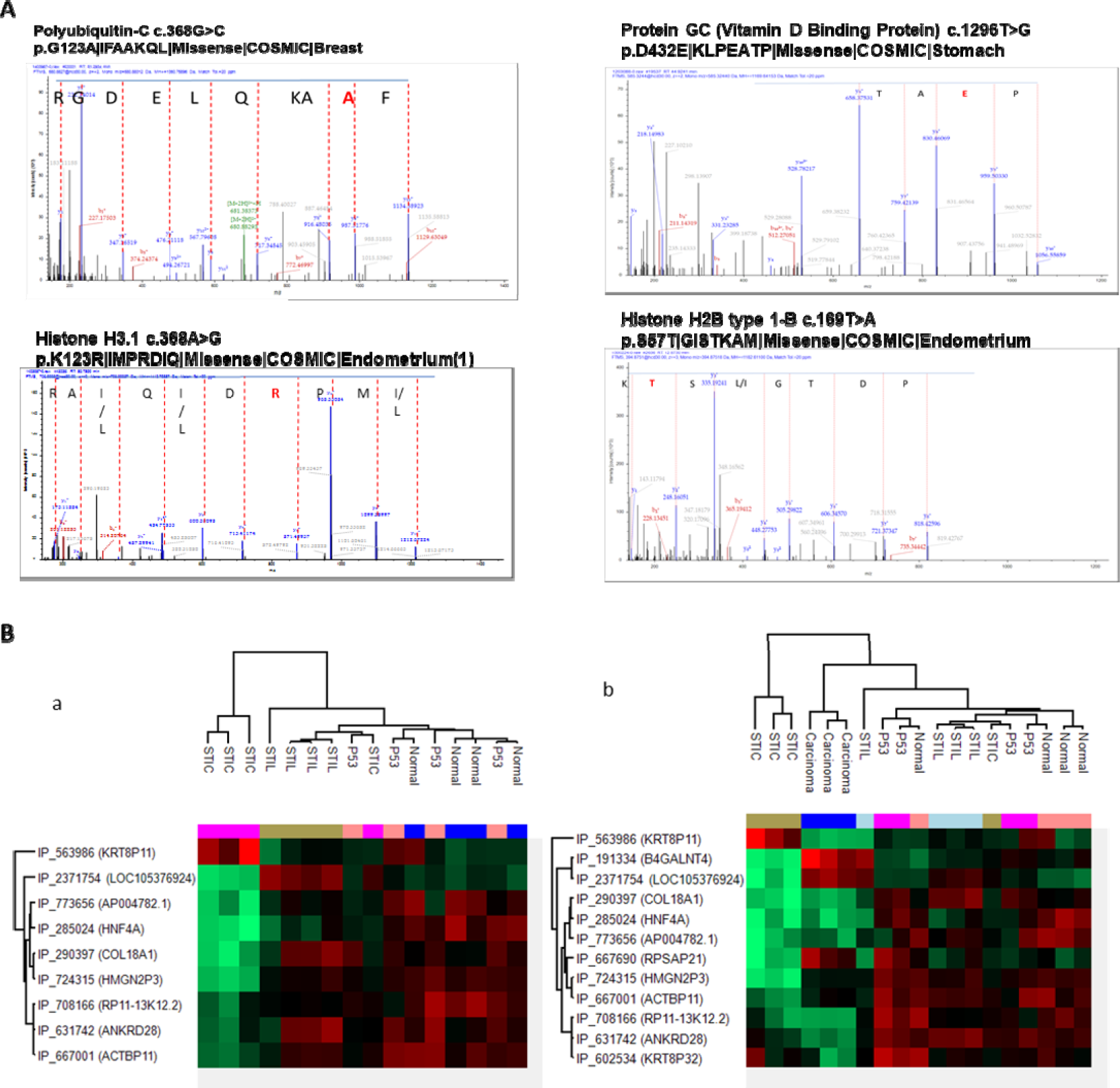
Potential mutations and Alternative proteins analysis. A) Annotated MS/MS spectra showing the mutated amino acid in red. B) Hierarchical clustering of the most variable alternative proteins between the lesion (a) or with adjunction of HGSC (b).

Considering the 83 mutations identified (**Table 3**, **Sup. Data 6**), some have been detected from normal to all lesion stages such as vimentin or collagen, but some seem specific of one particular transition *e;g.* mutations on Histone H4, Histone H2B type 1-C/E/F/G/I in p53 signature, for mutation on 60S ribosomal protein L14 in STIL, and mutation on (Na(+)/H(+) exchange regulatory cofactor NHE-RF1 in STIC. Interestingly, the transition from Normal to p53 signature, 7 mutated peptides have been identified. For p53 signature/STIL and Normal-p53/STIL, only one mutation has been observed respectively on Isoform 2 of Tropomyosin beta chain and Histone H1.4. From normal-p53/STIL and STIC, 3 mutated peptides have been found *e;g.* Laminin subunit gamma 1, Vimentin and Histone H3.1.and the last two mutations have been initially detected in endometrium.

**Table 3:** Mutation detected in proteins from proteomic data obtained by spatially resolved proteomic using XMAn database.

|  |  |  |  |  |
| --- | --- | --- | --- | --- |
| p53 signature | 2 | Histone H4 | c.239A>Gp.K80R\|AKRRTVT\|Missense | Endometrium |
|  | 2 | Histone H2B<br>type 1-C/E/F/G/I | c.181G>T p.G61C\|KAMCIMN\|Missense | Endometrium |
| STIL | 3 | 60S ribosomal<br>protein L14 | p.A159_K160insAA\|AAAAAKVP\|Insertion -<br>in frame | Endometrium,<br>Ovary, Breast |
| STIC | 2 | Na(+)/H(+)<br>exchange<br>regulatory<br>cofactor NHE-RF1 | c.767C>A p.P256H\|GPLHVPF\|Missense\| | Large intestine |
| Normal-p53<br>signature | 2/2 | Hemoglobin<br>subunit delta | c.427G>T p.A143S\|NALSHKY\|Missense | Lung |
|  | 2/2 | Histone H2B<br>type 1-C/E/F/G/I | c.133G>C p.V45L\|VYKLLKQ\|Missense | Breast |
|  | 2/2/1<br>HGSC | Actin, alpha<br>cardiac muscle 1 | c.166G>A p.V56I\|DSYIGDE\|Missense | Endometrium |
|  | 1/3 | Synaptic vesicle<br>membrane<br>protein VAT-1<br>homolog | c.278T>G p.L93R\|FADRMAR\|Missense | Lung |
|  | 3/3 | Isoform 2 of<br>Drebrin-like<br>protein | c.746G>A p.R249Q\|SRNQNEQ\|Missense | Endometrium |
|  | 2/3/1 STIL | Isoform 2 of<br>Synaptopodin-2 | c.3392C>T p.P1131L\|RPTLWEA\|Missense | Skin |
|  | 2/2/1 STIL | polyubiquitin c | c.368G>C p.G123A\|IFAAKQL\|Missense | Breast |
| p53signature/STIL | 2/3 | Isoform 2 of<br>Tropomyosin<br>beta chain | c.835C>A p.L279M\|QTLMELN\|Missense | Autonomic<br>ganglia |
| Normal-p53/STIL | 2/3/2:1<br>STIC | Histone H1.4 | c.262G>T p.V88L\|KSLLSKG\|Missense | Esophagus |
| Normal-p53/STIL/STIC | 1/3/3/2/1<br>HGSC | Vimentin | c.1313C>G p.S438*\|DTH\|Nonsense | Endometrium |
|  | 4/4/3/2 | Laminin subunit<br>gamma-1 | c.2033C>T p.P678L\|SARLGPG\|Missense | Breast |
|  | 4/4/4/4:<br>HGSC | Histone H3.1 | c.368A>G p.K123R\|IMPRDIQ\|Missense | Endometrium |

\*Only mutations found in more than two patients

### Ghost Proteome

Using our proteomic data against the OpenProt database, 59 AltProts have been identified. More than 64% of these proteins are associated to sequences from non-coding RNA (ncRNA), and from the mRNA coding for RefProt 23% are from 5’UTR, 33% from the 3’UTR and 42% from a shift in the CDS (**Supp. Data 7**). Among these proteins (**Table 4, Supp. Data 7**), IP_270787 (AltZNF709) is the single specific of normal tissue. IP_2326985 is issued from an ncRNA (LOC105376003) is only found in STIC. The AltProt IP_232994 (AltMYEF2) coding by the 3’UTR part of the transcript mRNA referenced to translate the MYEF2 is identified only in HGSC samples. IP_572777 (from the ncRNA: SDR16C6P), IP_774783 (from the ncRNA: RP11-358N4.6) and IP_285024 (from the ncRNA: HNF4A) are specific of Normal, p53 signature and STIL samples. In both Normal/p53/STIL and STIC samples, 17 AltProts were observed **(Table 4)**. Using the ANOVA test, 12 AltProts present a significant variation in the different lesion stages. Nine AltProt are derived from ncRNA, one from 3’UTR and two from a shift in the CDS. The hierarchical clustering and heatmap representation (**Figure 6Ba**) revealed a clear separation between STIC and the other lesion. The second branch separates STIL from normal-p53 signature. The pseudogene KRT8P11 (IP_563986) is overexpressed in STIC, whereas LOC105376924 and, COL18A1 (IP_290397) pseudogene are over-expressed in STIL. The other AltProts are differentially expressed in p53 lesions and Normal tissue, but all are under-expressed in STIC or non-expressed in STIL (**Figure 6Ba**). When data from HGSC is added, the separation of STIC from other samples is still observed, carcinoma samples composed the second branch, and a third branch is divided into subbranches. These subdivisions group three of the fourth STIL samples and three of the fourth Normal samples (**Figure 6Bb**). With regards to the results, KRTP8P11 (IP_563986) is a specific marker of STIC, B4GALN (IP_191334) is specific of HGSC and KRT8P32 (IP_602534) is overexpressed in two p53 signature samples and one of the normal tissue sample but also highly decreased in HGSC as ANKRD28 (IP_631742) and RP11-13K12.2 (IP_708166). COL18A1 (IP_290397) is specific to STIL. The other AltProt were observed between Normal-p53 signatures with most of them under-expressed in STIC (**Figure 6Bb**).

**Table 4:** Summary of the repartition to the Venn diagram result of the 45 AltProts identified. 14 AltProt have been deleted because they are identified but without a sufficient abundance to be quantified. The AltProt identification accession ID is according to the OpenProt Database. Venn diagram representation is based on the abundance obtain after analysis, function to the triplicate samples type (normal, p53, STIL, STIC, Carcinoma).

| Altprot | Normal | P53 | STIL | STIC | Carcinoma |
| --- | --- | --- | --- | --- | --- |
| IP_270787 | X |  |  |  |  |
| IP_572777 | X | X | X |  |  |
| IP_774783 | X | X | X |  |  |
| IP_285024 | X | X | X |  |  |
| IP_669028 | X | X | X | X |  |
| IP_563986 | X | X | X | X |  |
| IP_621829 | X | X | X | X |  |
| IP_595290 | X | X | X | X |  |
| IP_730500 | X | X | X | X |  |
| IP_2322038 | X | X | X | X |  |
| IP_763981 | X | X | X | X |  |
| IP_708166 | X | X | X | X |  |
| IP_737172 | X | X | X | X |  |
| IP_290397 | X | X | X | X |  |
| IP_652773 | X | X | X | X |  |
| IP_689518 | X | X | X | X |  |
| IP_721145 | X | X | X | X |  |
| IP_060976 | X | X | X | X |  |
| IP_632428 | X | X | X | X |  |
| IP_274301 | X | X | X | X |  |
| IP_138273 | X | X | X | X |  |
| **IP_716829** | **X** |  | **X** | **X** |  |
| **IP_2326985** |  |  |  | **X** |  |
| **IP_591310** |  | **X** | **X** | **X** | **X** |
| **IP_2285716** | **X** |  | **X** | **X** | **X** |
| **IP_572435** | **X** | **X** |  | **X** | **X** |
| **IP_2323408** | **X** |  |  | **X** | **X** |
| **IP_079312** |  | **X** | **X** |  | **X** |
| IP_667690 |  | X | X |  | X |
| IP_076499 |  | X | X |  | X |
| IP_601870 | X |  | X |  | X |
| IP_755940 | X | X |  |  | X |
| IP_232994 |  |  |  |  | X |
| IP_602534 | X | X | X | X | X |
| IP_667001 | X | X | X | X | X |
| IP_755869 | X | X | X | X | X |
| IP_658355 | X | X | X | X | X |
| IP_724315 | X | X | X | X | X |
| IP_591792 | X | X | X | X | X |
| IP_631742 | X | X | X | X | X |
| IP_688853 | X | X | X | X | X |
| IP_2371754 | X | X | X | X | X |
| IP_191334 | X | X | X | X | X |
| IP_773656 | X | X | X | X | X |
| IP_794359 | X | X | X | X | X |

## DISCUSSION

The most recent theories regarding the origin of HGSC propose an involvement of cells in the region of the tubal-peritoneal junction [66–69]. This junction corresponds to the region of the peritoneum covering the serosal surface of the fallopian tube and meets the specialized epithelium of the tubal fimbriae, with a transition between the Müllerian and the mesothelial cells [70]. Such junctional sites between different types of epithelia represent cancer hot spots in which neoplastic transformation occurs. This is confirmed by evidence that cells derived from this transition region demonstrate a cancer-prone stem cell phenotype [66, 68–70]. Subsequent studies led to the designation of these changes as STIC recognition of their preferential localization in the epithelium of the fimbriae end of the fallopian tube, and the recommendation of standardized and detailed sampling of tubal specimens with the SEE-FIM protocol. However, the diagnosis of STIC based on morphology alone is reported to have low reproducibility. With the emergence of detailed histological examination of the fallopian tubes, with and without IHC, a variety of confounding pre-neoplastic lesions is now being encountered which had not been previously characterized included p53 signature and STIL lesions [71], but the temporal relationship between each lesions and HGSCs is still debated.

The aim of our study was to contribute to the understanding the molecular mechanism associated with the development of the pre-neoplastic lesions found in the FTE. No similar proteomics studies have been published yet, because of the uniqueness of the protocol for sampling the tissue section. We firstly developed a novel strategy based on pathology routine protocol. Several tissue sections with pre-cancerous lesions is drastically limited. In fact, these types of lesions could only be observed using a SEE-FIM protocol as explained above. It allows localization and differentiation of each lesion type but on a limited number of slides. For example, p53 signature are observable only one or two tissue sections. In this context, the direct use of the IHC slide to perform other experiments has many advantages. In our study, to a better understanding of the different transitions of the lesions, *i.e*. from Normal to p53 signature, then from p53 signature to STIL and finally from STIL to STIC, IHC-guided spatially resolved proteomic analyses were realized on each identified lesion. We first demonstrated that the workflow we developed is compatible with proteomic analysis. Using targeted digestion of each region highlight by the IHC, we can preserve spatial information and perform analysis of different regions in the same slide. The tissue section surface recovery protocol does not alter the proteomic content and allows an accurate comparison of quantified proteins between each lesion. If we compare the proteomic content of HGSC and the lesion found in the fallopian tube, we observed interesting groupings that may correspond to what different studies propose for the chronology of HGSC development from pre-neoplastic lesions. The normal tissue and the p53 signature cluster together and are close to the STIL. The STIC lesion presents a more different signature and are closer to HGSC than the two other lesions. Due to the difference in composition between the fallopian tube tissue and the ovary, we have focused our analysis to deepen our understanding of the biological processes involved in the different pre-neoplastic lesions.

By comparing the global content of each lesion, we successfully identified specific markers for p53 signature, the protein RNA-binding protein 10 (RBM10)-3 specific proteins in STIC lesions -*i.e.,* Glucosamine (N-acetyl)-6-sulfatase (GNS), Upstream binding transcription factor, RNA polymerase I (UBTF) and ATPase family, AAA domain containing 3 (ATAD3A/B). We also identified specific mutated proteins from each lesion stages *i.e*. mutated Histone H4, Histone H2B type 1-C/E/F/G/I) in p53 signature, mutated 60S ribosomal protein L14 in STIL, and mutated (Na(+)/H(+) exchange regulatory cofactor NHERF1in STIC.

The p53 signature lesion is only characterized by a limited number of epithelial cells (∼10) presenting a p53 overexpression. In our analyses, the p53 signature samples are close in their protein content to the normal tissue. But some proteins specific to these lesions could be obtained. A signature emerged and demonstrated that among the 8 proteins overrepresented in p53 signature, some of them are unfavorable prognosis markers of OC (CAVIN1, SPTAN1 and FBLN5) and some for renal, head and neck cancers (EIF3B) or liver cancer (MOB1B,EMILIN-2, EIF3B), by contrast 3 are favorable prognosis markers for renal cancer (SPTAN1, SPTBN1, CAVIN2). We also dissected the molecular mechanisms that occur and the whole proteome analysis showed that the transition from normal to p53 signature was characterized by a residual immune response and translation activation, regulation of trafficking through cavins proteins, and ECM modifications through emilins.

From p53 signature to STIL: upregulated proteins are involved in the adhesion and anchoring of cells in the ECM, allowing a stabilization of the cells, along with an increase in maintenance activity instead of cell proliferation. Immune response and inflammation processes are still maintained, but an overall decrease of all metabolic processes, especially in TCA cycle is observed and with no Warburg effect occurs. These observations are in line with a recent study suggest that STIL could be considered as “Dormant STICs” and take a prolonged time of more than one decade to develop into STIC [17]. We observed that mostly pathway concerning metabolism and cell growth and maintenance is downregulated in STIL that could explain this “dormant” characteristic.

From STIL to STIC: Contrary to the dormant profile of STIL lesion, observations done on the different function dysregulation for STIC lesion, suggest, a more aggressive profile with a decrease of proteins involved in the cell adhesion and could reflect the start of an invasion step. Proteins up regulated in STIC show various functions inside the cells and appear to be involved in different steps of cancer processes. An increased of proteins involved in metabolism process and energy pathway are observed. If we compared the molecular functions involved in the three different groups, calcium-dependent protein binding is overrepresented. Also, heat shock protein activity is highly represented in this dataset. Proteins involved in telomere activities and centrosome maturation are specific to STICs compared to other lesions.

Interestingly, these two processes have been shown a link with early tumorigenesis [72] and that STIC precedes the development of many HGSCs [73]. A decrease of expression of proteins involved in the extracellular structure organization and cell adhesion molecule activity was observed in STIC lesion, suggesting a breaching of basement membrane and an increase of cell motility that could be conducted to an escape of these pre-cancerous cells to a distant organ. Finally, we observed a high level of proteins involved in extracellular vesicles including exosomes production which could consequently modify the microenvironment and promote cancerization processes.

From lesions to HGSC: One protein is found common in the pre-neoplastic lesions and HGSC, the Fibrillarin. FBL is one of the core proteins of box C/D small nucleolar ribinucleoprotein complexes (snoRNP). This complex is involved in first steps of pr-rRNA processing [74]. An overexpression of FBL contributes to tumorigenesis and confers cellular resistance to chemo drugs [75]. It has also been shown that a dysregulation of ribosome biogenesis plays key roles in oncogenesis [76]. Overactivation in cancer cells of ribosome biogenesis could be due to a loss of function of RNA polymerase repressors such as p53 [77]. Particularly FBL expression has been demonstrated to be correlated with p53 activity [78].

High levels of FBL protein were associated with the expression of mutant p53 and contributed to tumorigenesis by altering translational control of key cancer genes. UBTF is only present in STIC and HGSC. This protein is known to be a transcription factor involved in the regulation of the RNA polymerase I (Pol I), implicated in the regulation of cell cycle checkpoint and DNA damage response [79]and regulated by MYC [80]. Some studies show a link between Pol I initiation factor and chemoresistance of ovarian cancer [81]

Similarities between these pre-neoplastic lesions and HGSC have been found especially concerning *TP53* mutations and other genomic changes, but without excessive cell proliferation [82]. In that context, we decided to explore the presence of mutations at the protein level in the different steps of tubal tumor development. Using a specific database for protein mutation, XMAn, resulted in the identification of 83 mutated peptides sequences, of which 4 were observed directly in the tandem mass spectra. Taken together, it is interesting to observe that most of the protein mutated in p53 signature have been initially identified in endometrium cancer and are related to Histones and ribosome suggesting a link with the epigenetic and translation. Polyubiquitin C is related to proteasome and self-antigen presentation. Synaptopodin-2 and the Isoform 2 of Drebrin-like protein are actin-binding proteins known to be respectively an invasive cancer biomarker [83] and a potential marker for breast, lung and colorectal cancers [84]. Finally, the Actin, alpha cardiac muscle 1 is involved in cisplatin ovarian cancer cell resistance [85] and the synaptic vesicle membrane protein VAT-1 homolog is a marker of epithelial cells [86] and an oncogene in gastric cancer [87]. None of these mutated proteins have been found in STIL and STIC except the ones occurring on histones. Most of the mutated peptides observed in STIL, STIC, p53/STIL or in Normal/p53 signature/STIL/STIC are linked to the cytoskeleton and migration. These mutations could be key drivers of the cancer process from epigenetic modification step, and then on cytoskeleton modifications for cell migration and cancer development.

Using a dedicated database, identification of significantly variable AltProts, give us a new view of unknown markers. We previously established in HGSC and in glioma, the presence of cancer-promoting proteins from non-canonical human open reading frames (AltORF) [23–26, 88]. Among the alternative proteins (AltProt) observed in ovarian serous cancer, we already identified AltEDARADD, which now annotated by Uniprot with the accession number L0R849. Interestingly, a high expression level of AltEDARADD protein is associated with a poor prognosis for patients with ovarian HGSC [24]. Using Top-down proteomic strategy, we identified six AltProt specific to the benign region (AltTLR5, AltPKHD1L1, AtltSERPINE1, AltGPC5, AltGRAMD4, AltAGAP1), five in the necrotic/fibrotic tumor region (AltApol6, AltLARS2-AS1, AltLTB4R, AltTMP1, AltMTHFR) and four in the tumor (AltCMBL, AltGNL1, AltRP11-576E20.1, AltCSNK1A1L) [18]. AltGNL1 was selected for further analysis. In these conditions, AltGNL1 displayed a nuclear localization, whereas GNL1 was present in the cytosol [18]. In our present study, some of the identified AltProt or transcripts have been described in other cancer studies. The pseudogene KRT8P32 has been identified in breast cancer [89], the pseudogene ACTBP11 is observed in GM12878 (B cells) cell line according to Diana-LncBase V3 [90]. Concerning the ANKRD28, this AltProt has been recently demonstrated to act as a novel BRCA1-interacting protein in breast and ovarian cancer [91]. RP11-13K12.2 is described to be overexpressed in pediatric ovarian fibrosarcoma [92]. COL18A1 and B4GALNT4 are overexpressed in endometrial and ovarian cancers according to the HPA and are unfavorable markers for overall survival for both cancers. Some of these are specific to the type of lesion - *i.e.* LOC105376924 and COL18A1 in STILL, KRT8P11 in STIC and ANKRD28 in STILL-STIC. These alternative proteins seem to be clinically relevant, and further exploration will lead to a better understanding of the disease and new targets to improve diagnosis.

Finally, the Identification of the AltProt and the high number of mutations affecting histones and cytoskeleton proteins observed between normal to p53 signature compared to the other lesions, are in line with the hypothesis of epigenetic reprogramming towards carcinogenesis and cell transformation [93]. A recent study established that epigenetic reprogramming occurs specifically in the proximal end of the fallopian tubes in *BRCA* mutation carriers. This epigenetic reprogramming event is driven by aberrantly high AICDA (also named AID, activation-induced cytosine deaminase) expression and is an integral early pre-malignant event in HGSC development. In this context, our result would provide an interesting starting point for further studies, particularly with respect to the potential link to endometrium and exfoliation of endometrial carcinoma cells due to epigenetic reprogramming to carcinogenesis [4] or that STIL represents exfoliated precursor cells that eventually undergo malignant transformation within the peritoneal cavity [94].

Overall, the present study identified several processes involved in the various neoplastic lesions that are observed in the fimbriated end of the fallopian tube. Besides, we have demonstrated the association across the different lesions, which may be useful in identifying potential timeline and underlying mechanisms of origin of ovarian cancer.

## Supporting information

Abstract

## ACKNOWLEDGEMENTS

This research was supported by grants from the Ministère de L’Education Nationale, de L’Enseignement Supérieur et de la Recherche, ANR (IF), SIRIC ONCOLille (IF), Grant INCa-DGOS-Inserm 6041aa, and INSERM, the PHRC FIMBRIA (EL), Ligue contre le Cancer (MW).

## Author contributions

IF, MW and MS conceived and supervised the study. MW, PS performed the experiment. Surgery was realized by EL and FN. The pathology analyses were done by YMR DB, ASL. FK, SA, MW performed the systemic biology analyses. IF, EL, MW, PS and MS participated in experimental design. MW, SA and PS performed data analyses. MS and MW wrote the manuscript with contributions from all co-authors. MS, EL, MW and IF found funds for the project.

## Additional Information

The authors declare no competing financial interests in this work.

## Abbreviations list

FTE: fallopian tube epithelium
EMT: Epithelial-mesenchymal transition
FFPE: formalin fixed paraffin embedded GO Gene Ontology
HGSC: High Grade Serous Carcinoma
HPA: Human Protein Atlas
IHC: immunohistochemistry
MALDI: Matrix assisted Laser Desorption Ionization
MSI: Mass spectrometry imaging
OC: Ovarian cancer OS overall survival
PCA: principal component analysis
SEE-FIM: Sectioning and Extensively Examining the Fimbriated End Protocol
STIC: serous tubal intraepithelial carcinoma
STIL: serous tubal intraepithelial lesions
TF: Transcription factor

## SUPPORTING INFORMATION

Main Text: 6 Figures and 4 Tables.

Supplementary files:

Supp. Data 1: List of quantified proteins corresponding to Venn diagram for lesions comparison

Supp. Data 2: List of quantified proteins corresponding to Venn diagram for lesions and HGSC comparison

Supp. Data 3: List of quantified proteins significantly modified in the different preneoplastic lesions

Supp. Data 4: Significantly enriched pathways obtained by enrichment analysis using Panther Db

Supp. Data 5: Quantitative comparison of proteins modified between normal tissue and p53 signature

Supp. Data 6: List of mutated peptides identified using the XMAn database

Supp. Data 7: List of alternatives proteins identified

Supplementary figures:

**Figure S1:**
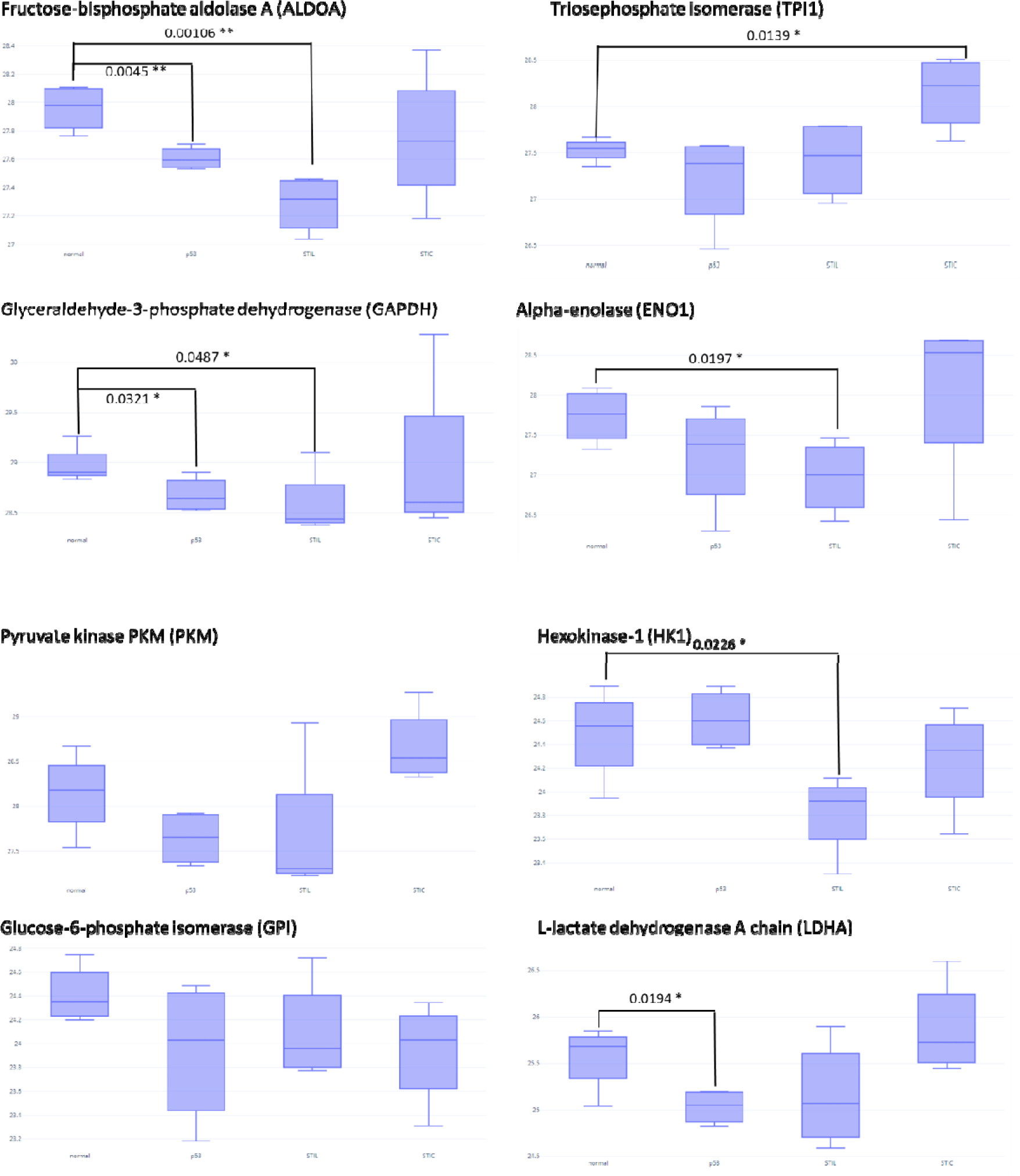
Visualization of protein levels in the different lesions for proteins known to be involved in the Warburg effect. Log(LFQ) values were used for the representation of Fructose-bisphosphate aldolase A (ALDOA), Triosephosphate isomerase (TPI1), Glyceraldehyde-3-phosphate dehydrogenase (GAPDH), Alpha-enolase (ENO1), Pyruvate kinase PKM (PKM), Hexokinase-1 (HK1), Glucose-6-phosphate isomerase (GPI) and L-lactate dehydrogenase A chain (LDHA). A t-test is used to compare the value between normal tissue and each lesion. P-value is represented by stars (*** p < 0.001, ** p < 0.01, * p < 0.05 no star for p > 0.05).

